# PHENSIM: Phenotype Simulator

**DOI:** 10.1101/2020.01.20.912279

**Authors:** Salvatore Alaimo, Rosaria Valentina Rapicavoli, Gioacchino P. Marceca, Alessandro La Ferlita, Oksana B. Serebrennikova, Philip N. Tsichlis, Bud Mishra, Alfredo Pulvirenti, Alfredo Ferro

## Abstract

Despite the unprecedented growth in our understanding of cell biology, it still remains challenging to connect it to experimental data obtained with cells and tissues’ physiopathological status under precise circumstances. This knowledge gap often results in difficulties in designing validation experiments, which are usually labor-intensive, expensive to perform, and hard to interpret.

Here we propose PHENSIM, a computational tool using a systems biology approach in order to simulate how cell phenotypes are affected by the activation/inhibition of one or multiple biomolecules and does so by exploiting signaling pathways. Our tool’s applications include predicting the outcome of drug administration, knockdown experiments, gene transduction, and exposure to exosomal cargo. Importantly, PHENSIM enables the user to make inferences on well-defined cell lines and includes pathway maps from three different model organisms. To assess our approach’s reliability, we built a benchmark from transcriptomics data gathered from NCBI GEO and performed four case studies on known biological experiments. Our results show high prediction accuracy, thus highlighting the capabilities of this methodology.

PHENSIM standalone Java application is available at https://github.com/alaimos/phensim, along with all data and source codes for benchmarking. A web-based user interface is accessible at https://phensim.atlas.dmi.unict.it/.

## Introduction

Cells of living organisms are continuously exposed to signals originating in both the extracellular and the intracellular microenvironments. These signals regulate multiple cellular functions, including gene expression, chromatin remodeling, DNA replication and repair, protein synthesis, and metabolism. The proper response to signals depends on the expression, activation, or inhibition of sets of interrelated genes/proteins, acting in a well-defined order within the framework of vector-driven biological processes, aiming to reach specific endpoints. Such sub-cellular processes are referred to as biological pathways [1].

In this context, the study of genome and transcriptome, the definition of protein-protein interaction networks, and association studies between gene sets and molecular mechanisms in humans have produced valuable biological information. However, despite the improvements in our understanding of cell biology, it is challenging to link omics data to the physiopathological status of cells, tissues, or organs under specific conditions. Besides, studies addressing these issues are often labor-intensive, expensive to perform and produce big datasets for analysis.

Recently, systems biology computational approaches have emerged as efficient means capable of bridging the gap between experimental biology at the system-level and quantitative sciences [2]. Indeed, such methods can be used as time- and cost-saving solutions for efficient *in silico* predictions [2, 3]. Here, network analysis is playing a central role by modeling and understanding of biological phenomena. In this perspective, simulation methodologies can help understand the intricate interaction patterns between molecular entities, significantly improving manual analysis. Furthermore, ‘in silico’ simulations can be extensively applied in massive-scales, testing thousands of hypotheses under various conditions, which is usually experimentally infeasible.

At present, a large number of simulation models have become available. However, they can be grouped into two broad categories: (i) discrete/logic, or (ii) continuous models [4]. Discrete models represent each element’s state in a biological network as discrete levels, and the temporal dynamic is also discretized. At each time step, the state is updated according to a function, determining how an entity’s state depends on the state of other (usually connected) entities. Boolean networks [5, 6] and Petri nets [7] represent two types of discrete models. BioNSi (Biological Network Simulator) [8] is an intuitive model, implemented as a Cytoscape 3 plugin [9]. It can use KEGG pathways [10] as a network model and represents each element in discrete states (usually up to 10). At each simulation time point, the state of a node is updated using an effect function. The simulation ends as soon as it reaches a steady-state. The model is easy to use. However, a more complex biological network might pose challenges to its performance.

Continuous models usually produce real continuous measurements, instead of discretized values, simulating network dynamics over a continuous timescale. Although they could provide a greater degree of accuracy, these methods are limited by our current description of the biological systems and our measurement techniques’ capabilities. Continuous linear models [11, 12], and flux balance analysis [13] are some of the most representative continuous models.

Here, we present PHENSIM (PHENotype SIMulator), a web-based, user-friendly tool allowing phenotype predictions on selected cell lines or tissues in three different organisms: *Homo sapiens, Mus musculus*, and *Rattus norvegicus*. PHENSIM uses a probabilistic algorithm to compute the effect of dysregulated genes, proteins, microRNAs (miRNAs), and metabolites on KEGG pathways. Results are summarized through a Perturbation, which represents the expected magnitude of the alteration, and an Activity Score, which is an index of both the predicted effect of a gene dysregulation on a node (up- or down-regulation) and its likelihood. All values are also computed at the pathway-level. Moreover, to achieve greater accuracy, PHENSIM performs all calculations in the KEGG meta-pathway, obtained by merging all pathways [14] (see Methods) and integrates information on miRNA-target and transcription factor (TF)-miRNA extracted from online public knowledge bases [15]. We implemented our tool as a freely accessible web application at the following URL: https://phensim.atlas.dmi.unict.it/

## Results

To assess the performances of PHENSIM, we performed a comprehensive experimental analysis. As detailed in the section “Experimental Procedure and Benchmarking”, we built a benchmark, ran four different simulations as case studies and manually analyzed their results.

The benchmark was built by taking public GEO series of up-/down-regulation of single genes in cell lines. We acquired 22 series further divided into 50 case/control sets. The sets were categorized based on the genes present in the KEGG meta-pathway (DS1 contains all sample sets where the up- or down-regulated gene was in KEGG; DS2 all the other samples). PHENSIM and BioNSi simulations were evaluated in terms of Accuracy, Positive Predictive Value (PPV), Sensitivity and Specificity for genes showing altered expression, and PPV and False Negative Rate (FNR) for the others.

Our results show that PHENSIM has an average accuracy of 0.6647 for the dataset in the first category and 0.5977 for the second category. Whereas BioNSi offers an average accuracy of 0.0640 and 0.0735 for the datasets in the first and second categories. Nevertheless, PHENSIM has comparable PPV to BioNSi (0.5123 and 0.5075, respectively) in the first category, while a higher PPV can be observed for the second category (PHENSIM=0.7725, BioNSi=0.3282). PHENSIM also shows a greater Sensitivity and Specificity to BioNSi. Supplementary Table S1 reports the detailed comparison in terms of average metrics.

To assess performance differences between the two systems for each dataset, we provide several graphs comparing each metric. In Figure 1, we summarize the results from the DS1 datasets, and in Figure 2, we report the results from the DS2 datasets. In each graph, we detail a single metric: Positive Predictive Value (PPV), Sensitivity and Specificity for genes showing altered expression, and PPV and False Negative Rate (FNR) for the non-altered ones. On the x-axis, we have PHENSIM performance, while on the y-axis, we have BioNSi. Each dot represents a dataset. The black line marks the points where the two algorithms have the same performance.

**Figure 1.**
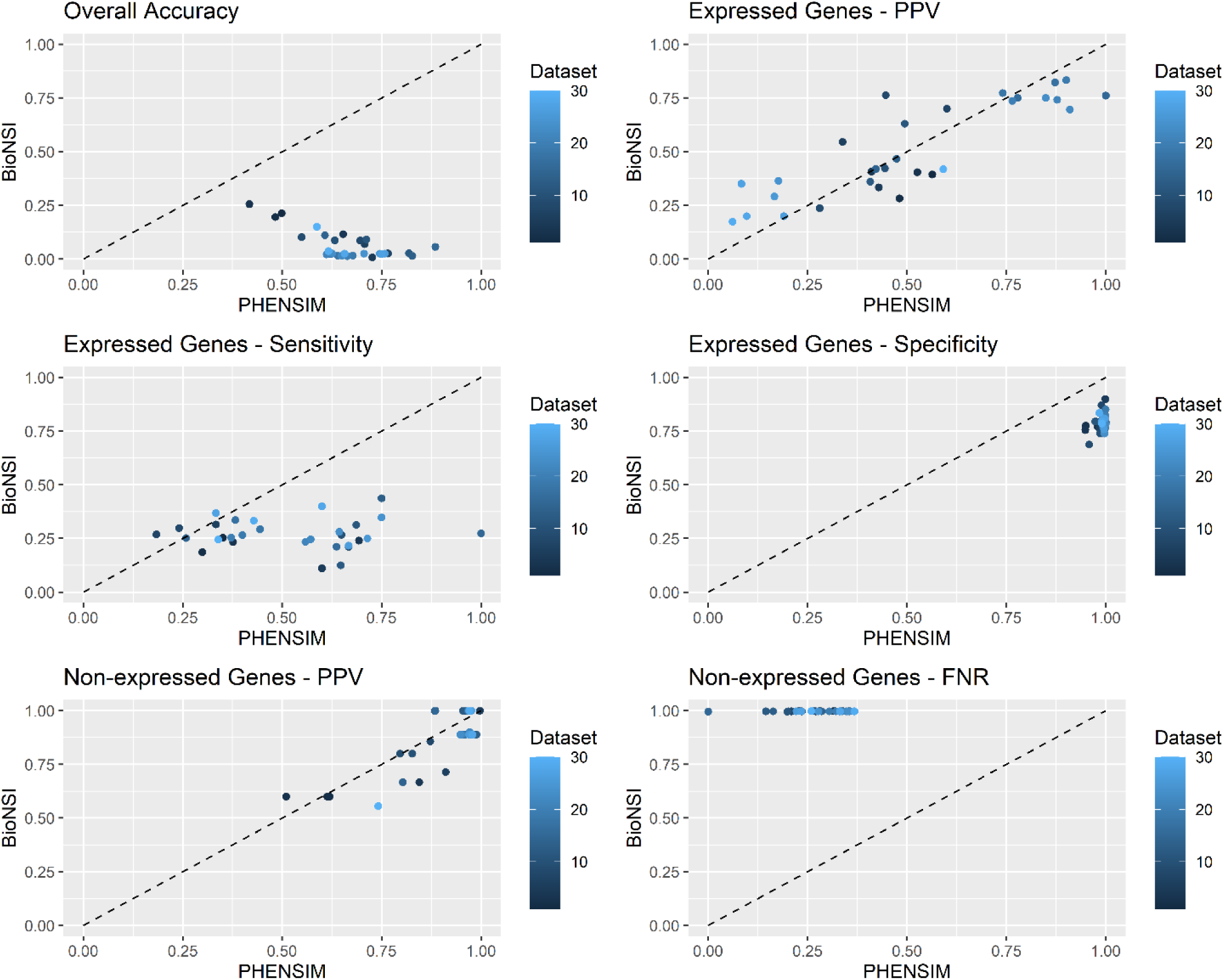
Comparison between PHENSIM and BioNSi for datasets where the altered gene was in the meta-pathway. Each graph reports one metric: Positive Predictive Value (PPV), Sensitivity and Specificity for genes showing altered expression, and PPV and False Negative Rate (FNR) for the non-altered ones. On the x-axis, we report PHENSIM performance while on the y-axis, we present BioNSi. Each dot represents a dataset. The black line marks the points where the two algorithms have the same performance. On a dataset below the line, PHENSIM has better performance than BioNSi; above the line it is the opposite.

**Figure 2.**
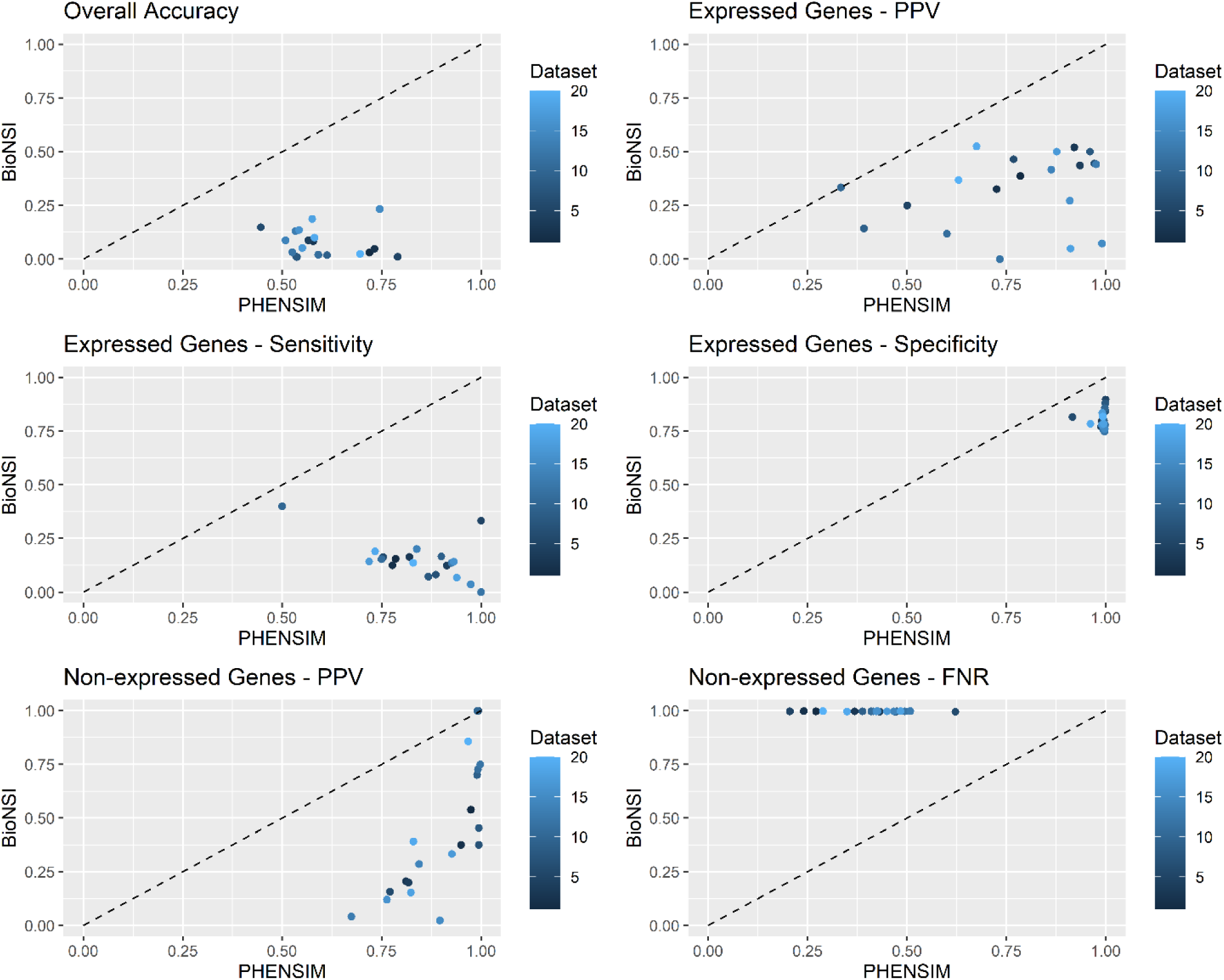
Comparison between PHENSIM and BioNSi for datasets where the altered gene was not in the meta-pathway. Each graph reports one metric: Positive Predictive Value (PPV), Sensitivity and Specificity for genes showing altered expression, and PPV and False Negative Rate (FNR) for the non-altered ones. On the x-axis, we report the PHENSIM performance while on the y-axis, we have BioNSi. Each dot represents a dataset. The black line marks the points where the two algorithms have the same performance. On a dataset below the line, PHENSIM has better performance than BioNSi; above the line it is the opposite.

Finally, to complete our assessment of PHENSIM capabilities, we run several simulations to perform 4 case studies on known biological experiments: (i) anti-cancer effects of metformin, (ii) Everolimus (RAD001) treatment in breast cancer, (iii) effects of exosomal vesicles on hematopoietic stem/progenitor cells (HSPCs) in the bone marrow (BM) and (iv) testing TNFα/siTPL2-dependent synthetic lethality on a subset of human cancer cell lines. We examined the ability of PHENSIM to correctly predict the activity status of both individual genes/proteins and signaling pathways by comparing PHENSIM predictions with experimental data. In the following sections, we briefly report the results of two case studies: the anti-cancer effects of metformin and the testing TNFα/siTPL2-dependent synthetic lethality on a subset of human cancer cell lines. Detailed descriptions of all case studies are provided in the Supplementary Materials.

### Anti-cancer effects of metformin

Metformin is a widely prescribed agent for the treatment of type 2 diabetes [16-19]. It inhibits glucose production in the liver and increases insulin sensitivity in the peripheral tissues. Furthermore, metformin treatment reduces insulin secretion by β-pancreatic cells. The key molecule that executes these functions is AMP-activated protein kinase (AMPK). Several evidence indicates that metformin may also possess anti-cancer effects, especially in diabetic patients [16-18]. One of its major drivers seems to be the LBK1-AMPK signaling pathway [18]. An overview of the metformin-mediated effects is reported in Figure 3.

**Figure 3.**
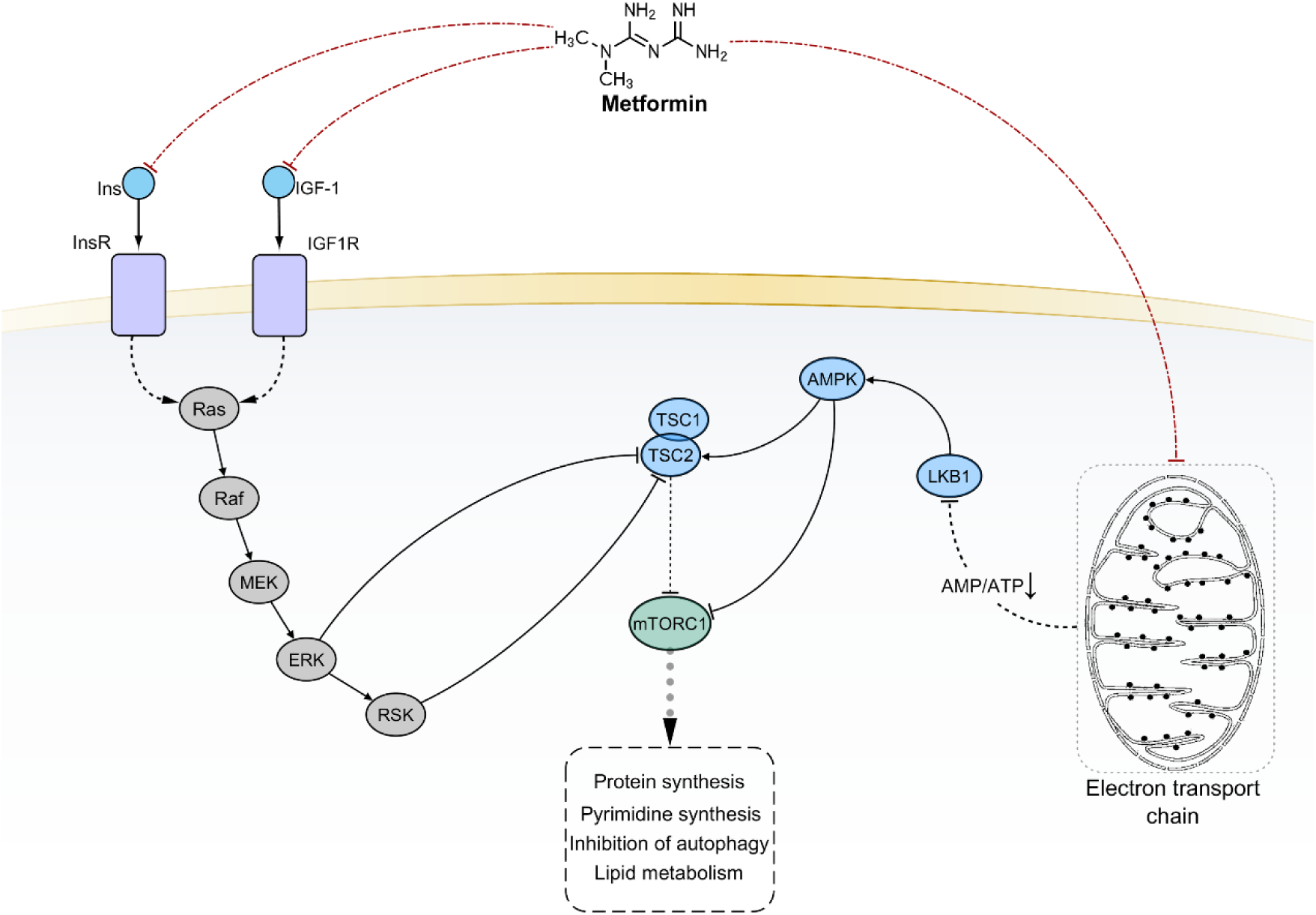
The current model of metformin-mediated pharmacological effects. Black solid edges represent direct interaction between first neighbor nodes. Dashed edges represent indirect interactions between nodes. Red dot-dashed edges evidence scientifically validated interactions considered for PHENSIM prediction.

We ran PHENSIM to simulate the simultaneous upregulation of LKB1 and the downregulation of both insulin (Ins), IGF1, and GPD1 [20]. As expected, PHENSIM returned significant downregulation of Insulin and mTOR signaling (Insulin activity score = -8.7121, p-value 0.072; mTOR activity score = -8.7121, p-value 0.075). PI3K (phosphoinositide 3-kinase), AKT (serine/threonine protein kinase Akt), and metabolite PIP3 (phosphatidylinositol (3,4,5)-trisphosphate) were also downregulated (PIK3CA activity score = -4.8203, p-value 0.021; AKT1 activity score = -4.2546, p-value 0.04). We also predicted the negative regulation of mTOR (activity score = -3.9992; p-value = 0.04), though no prediction were made for the *repressor of translation initiation* 4EBP. However, a low positive perturbation was detected. PHENSIM also predicted the inhibition of downstream nodes involved in protein synthesis as S6Ks (fig. S1a). Furthermore, a low perturbation for TCA cycle (perturbation = -0.0000177) can be observed.

MAPK (activity score = -8.7121, p-value 0.074) and NF-*κ*B (NF-*κ*B activity score = -4.8203, p-value 0.111) signaling were predicted downregulated. Furthermore, several downregulated enzymes and metabolites were correctly detected by PHENSIM (fig. S1b).

### Testing TNFα/siTPL2-dependent synthetic lethality on a subset of human cancer cell lines

TNFα (tumor necrosis factor alpha), a type II transmembrane protein, is a member of the tumor necrosis factor cytokine superfamily and has an essential role in innate immunity and inflammation. Although it can induce cell death, most cells are protected by a variety of rescue mechanisms.

In a recent paper, Serebrennikova et al. [21] showed that TPL2 (MAP3K8) is one of the TNFα-induced cell death checkpoints. Its knockdown resulted in the downregulation of miR-21 and the upregulation of its target CASP8 (caspase-8). This response, combined with the downregulation of caspase-8 inhibitor cFLIP (FADD-like IL-1β-converting enzyme inhibitory protein), resulted in the activation of caspase-8 by TNFα and the initiation of apoptosis (Fig. 4). The activation of caspase-8 also promotes the activation of the mitochondrial pathway of apoptosis. It is worth noticing that the activation of the apoptotic (caspase-8-dependent) pathway in TNFα/siTPL2 treated cells was observed in some, but not all cancer cell lines, suggesting that correct prediction will depend on whether the data analyzed by PHENSIM are derived from sensitive or resistant cells.

**Figure 4.**
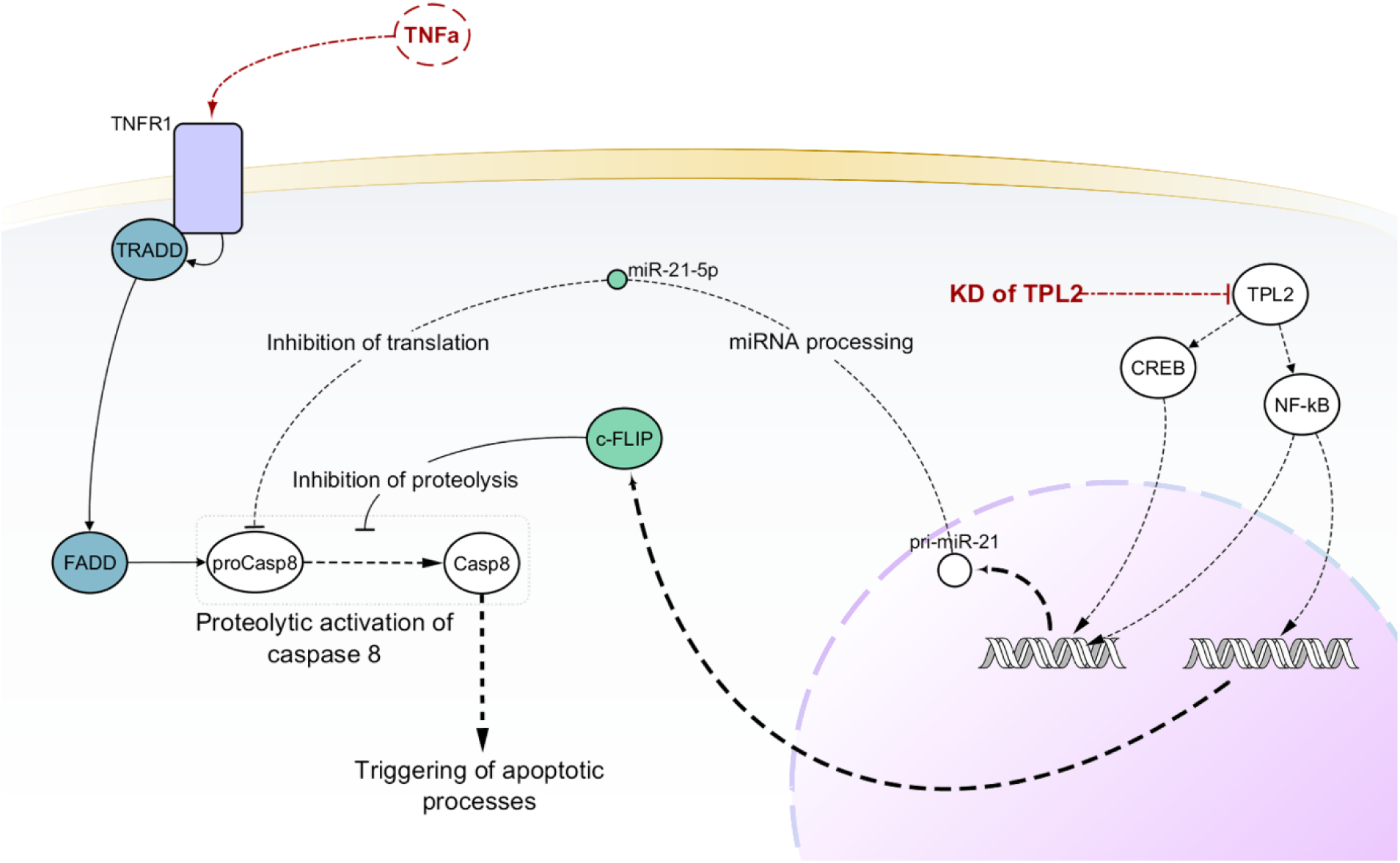
Generalized model showing molecular mechanisms underlying the TNFα/siTPL2-dependent synthetic lethality. Black solid edges represent direct interaction between first neighbor nodes. Dashed edges represent indirect interactions between nodes. Red dot-dashed edges evidence scientifically validated interactions considered for PHENSIM prediction.

To start the simulation, we set TPL2 and miRNA-21-5p as downregulated and TNFα as up-regulated. Since our goal was to simulate the outcome of such treatment in six different cell lines (HeLa, HCT116, U2-OS, CaCo-2, RKO, and SW480), we ran six simulations. Each simulation had a diverse list of non-expressed genes, one for each cell-line.

Among these tumor cell lines, only HeLa, HCT116, U2-OS were sensitive to treatment with TNFα/siTPL2. PHENSIM was able to correctly predict the upregulation of caspase-8 for the six cell lines. However, PHENSIM could not detect the downregulation of cFLIP in the resistant cell lines. PHENSIM did not predict any activity score for MCL1 (Mcl-1 apoptosis regulator) and XIAP (X-linked inhibitor of apoptosis). However, a low negative perturbation can be observed in all cell lines except HCT116 (where it is positive).

PHENSIM predicted the upregulation of the apoptosis inhibitors BCL2 and BCL-XL in all sensitive cell lines (only for HCT116, the activity score is 0, but the perturbation is still positive). BCL2 is also up-regulated in RKO and SW480 cells. PHENSIM showed a negative perturbation of the inducer of mitochondrial apoptosis BAX only in U2-OS and HELA among the sensitives, and SW480 and SW480 as resistant cell lines. (fig. S6a-d).

Although these results do not entirely reflect our expectations as there are discrepancies between the in vitro experiment and our predictions, it was confirmed by results obtained in Serebrennikova et al. [21] that the change in the expression of such molecules was due to the activation of feedback mechanisms. Interestingly, this result was obtained only for four out of six cancer cell lines, of which three were sensitive (HeLa, HCT116, and U2-OS), and one was resistant (CaCo-2).

Furthermore, phosphorylated ERK, MEK, JNK, and p38 activity were strongly downregulated for all cell lines. Finally, cIAP2 (baculoviral IAP repeat-containing 2) had activity score 0 and a weak negative perturbation in HTC116, HeLa, U2OS, SW480, and CaCo-2 cells, as confirmed by the experimental data (fig S6a-b).

## Discussion

This paper introduces PHENSIM, a flexible, user-friendly pathway-based simulation technique and an in silico tool based on it. PHENSIM has been mainly developed to predict the effects of one or multiple molecular deregulations on cell/tissue phenotype. Thus, we view PHENSIM as an easy-to-use, supportive pathway-based method that can make predictions of in vitro experiments targeting the expression of signaling processes’ activity.

To evaluate our tool’s potential, we built a benchmark of 50 case/control sample sets derived from 22 GEO series. Each set contained expression data of experiments regarding the up- or down-regulation of one single gene in a specific cell line. As previously described, 30 sample sets were directly used since the tested gene was already in KEGG. The remaining 20 sets were simulated through their differentially expressed genes (DEGs). Here, the main idea is that the DEGs can summarize the downstream alterations caused by the experiment. We compared our approach’s performance with BioNSi, a Cytoscape plugin for modeling biological networks and simulating their dynamics. Results-based comparative evaluations were performed in terms of accuracy, Positive Predictive Value (PPV), Sensitivity and Specificity for genes showing altered expression, and PPV and False Negative Rate (FNR) for the non-altered ones.

We show that, on average, our tool obtains better results than BioNSi in terms of accuracy, PPV, Sensitivity, Specificity, and FNR. More in detail, for the 30 samples of DS1, we show that only in 11 cases BioNSi achieves a greater PPV than PHENSIM. However, Sensitivity and Specificity are still higher for our methodology. In the other 20 samples, PHENSIM always outperforms BioNSi. Furthermore, when looking at non-expressed genes, BioNSi has significantly higher FNR than PHENSIM.

To further explore PHENSIM capabilities, we performed four case studies in different scenarios: drug administration to cultured cells (simulations 1 and 2), effects of exosomal-derived miRNAs in recipient cells (simulation 3), and the combined targeting of two signaling molecules, which are known to induce synthetic lethality in a subset of cell lines (simulation 4). After comparison, the literature data and PHENSIM predictions were in almost full agreement with simulation #1 and in partial agreement with the three remaining simulations, showing a discrete degree of accuracy.

Discrepancies with baseline data sugest some limitations in the predictive potential of our method. However, since pathway analysis relies on prior knowledge about how genes, proteins, and metabolites interact, we hypothesize that such a negative outcome is at least partly due to the incompleteness of the existing knowledge employed in the study. Indeed, since the biological pathways on current databases are still largely fragmented, calculations based on them will inevitably produce less than ideal results [22]. One example of this limitation is provided by mTORC1 downstream signaling. It is known that mTORC1 promotes protein synthesis by phosphorylating p70SK and 4EBP. It also stimulates ribosome biogenesis via inhibitory phosphorylation of the RNA Polymerase III repressor MAF1 [23]. mTORC1-induced pyrimidine biosynthesis is stimulated by p70S6K-mediated phosphorylation of the CAD enzyme (carbamoyl-phosphate synthetase 2, aspartate transcarbamylase, and dihydroorotase). Furthermore, the upregulation of 5-phosphoribosyl-1 pyrophosphate (PRPP) is an allosteric activator of CAD [24, 25].

KEGG Pathways do not consider such interactions. Therefore, our tool could not predict any perturbations for these biological processes. Similar observations can be made for the downregulation of cFLIP in the siTPL2/TNFα-resistant cell lines by our method. However, we were able to identify indirect evidence of such activity. On the other hand, the correct predictions obtained for autophagy, RNA transport, and mTOR signaling in simulation 2, and the mitochondrial apoptotic pathway activation in simulation 4, suggest that, provided with the right information, PHENSIM is likely to obtain significantly better results.

A further limitation for pathway analysis methods is the current knowledge-bases’ inability to contextualize gene expression and pathway activation in a cell- and condition-specific manner [22]. Furthermore, pathways do not consider protein isoforms encoded by different genes or differently processed mRNAs derived from a single gene. This poses a significant limitation since such isoforms may have unique and sometimes opposite signaling properties. By developing a strategy that allows removing non-expressed genes from the computation, we offer the user the possibility to contextualize predictions in a cell- or tissue-dependent manner. In conjunction with this, the integration of KEGG pathways with information coming from post-transcriptional regulators such as miRNAs increased the accuracy of the results, thus leading to considerable improvements of predictions [26]. Moreover, using the KEGG meta-pathway approach, instead of single disjointed pathways, partially addresses pathway independence [22].

In conclusion, PHENSIM showed good accuracy in most applications and could predict the effects of several biological events starting from the analysis of their impact on KEGG. We believe that several discrepancies can be traced to the incompleteness of knowledge in KEGG pathways or the lack of appropriate cell- and condition-specific information. Such incompletenesses can be partially addressed through a manual annotation of the pathways with the missing elements and links, including miRNA-target and TF-miRNA interactions. Furthermore, we plan to add other pathway databases such as Reactome or NCI pathway to enhance our meta-pathway. PHENSIM is limited to the simulation of changes in the expression or activity of signaling molecules. It is not suitable to simulate genetic aberrations unless they affect molecules expression or activity directly. Despite these limitations, our approach shows appreciable utility in the experimental field as a tool for the reliable prioritization of experiments with greater success chances.

## Methods

### Overview of the method

PHENSIM is a randomized algorithm to predict the effect of (up/down) de-regulated genes, metabolites, or microRNAs on the KEGG meta-pathway [14]. The meta-pathway is a network obtained by merging all KEGG pathways through their common nodes. This approach allows us to consider pathway crosstalk and, ideally, gives a more comprehensive representation of the human cell environment. Furthermore, the KEGG meta-pathway is annotated with experimentally validated miRNA-target and Transcription Factor-miRNA interactions to consider post-transcriptional expression modulation.

Currently, our method uses all KEGG pathways (downloaded on April 2020) with details on validated miRNA-targets inhibitory interactions downloaded from miRTarBase (release 8.0) [27] and miRecords (updated to April 2013) [28], and TF-miRNAs interactions obtained from TransmiR (release 2.0) [29].

To start a simulation, PHENSIM requires a set of nodes (at least one) together with their “deregulation type” (up-/down-regulation) as input values. We can also provide: (i) a list of non-expressed genes, (ii) a set of new nodes or edges that will be added to the KEGG meta-pathway, and (iii) the organism (*H. sapiens, M. musculus*, and *R. norvegicus)*. For the sake of clarity, we first define the case when input elements are independently altered. That is, input nodes whose expression is independently changed from one another (i.e., transfection of two siRNAs for knockdown of two genes). Next, we report an efficient and reliable technique to deal with dependent alterations.

PHENSIM uses the input to compute synthetic Log-Fold-Changes (LogFC) values. These values are then propagated within biological pathways using the MITHrIL algorithm proposed in Alaimo et al. 2016 [15], to establish how these local perturbations can affect the cellular environment. This propagation result is called a “Perturbation”, and it reflects the change of expression for a gene in a pathway (negative/positive for down-/up-regulation). This value is computed for each gene in the meta-pathway. Finally, PHENSIM summarizes all results using two values for each gene: the “*Average Perturbation*” and the “*Activity Score*” (*AS*). The average perturbation is the mean for all perturbation values computed during the simulation process and reproduces the expected change of expression for the entire process. The function of the Activity Score is twofold. The sign gives the type of predicted effect: positive for activation, negative for inhibition. The value is the log-likelihood that this effect will occur. Together with the AS, PHENSIM also computes a p-value through a bootstrapping procedure. All p-values are then corrected for multiple hypotheses using the q-value approach [30]. PHENSIM p-values are used to establish how biologically relevant the predicted alteration is for the simulated phenomena - i.e., the lower is a node p-value, the less likely it is that such alteration will occur by chance. An overview of the PHENSIM algorithm is depicted in Figure 5. The algorithm comprises of 5 main steps. Given a user input, (i) synthetic LogFC are generated and a (ii) simulation step is performed. These steps are repeated 1000 times to (iii) compute the *AS*. Next, user input is (iv) randomized, and 100 synthetic LogFC are generated to estimate *AS* using the simulation step. The input is randomized 1000 times to obtain greater precision. Finally, (v) p-values are computed, and the False Discovery Rate is estimated using the q-value methodology.

**Figure 5.**
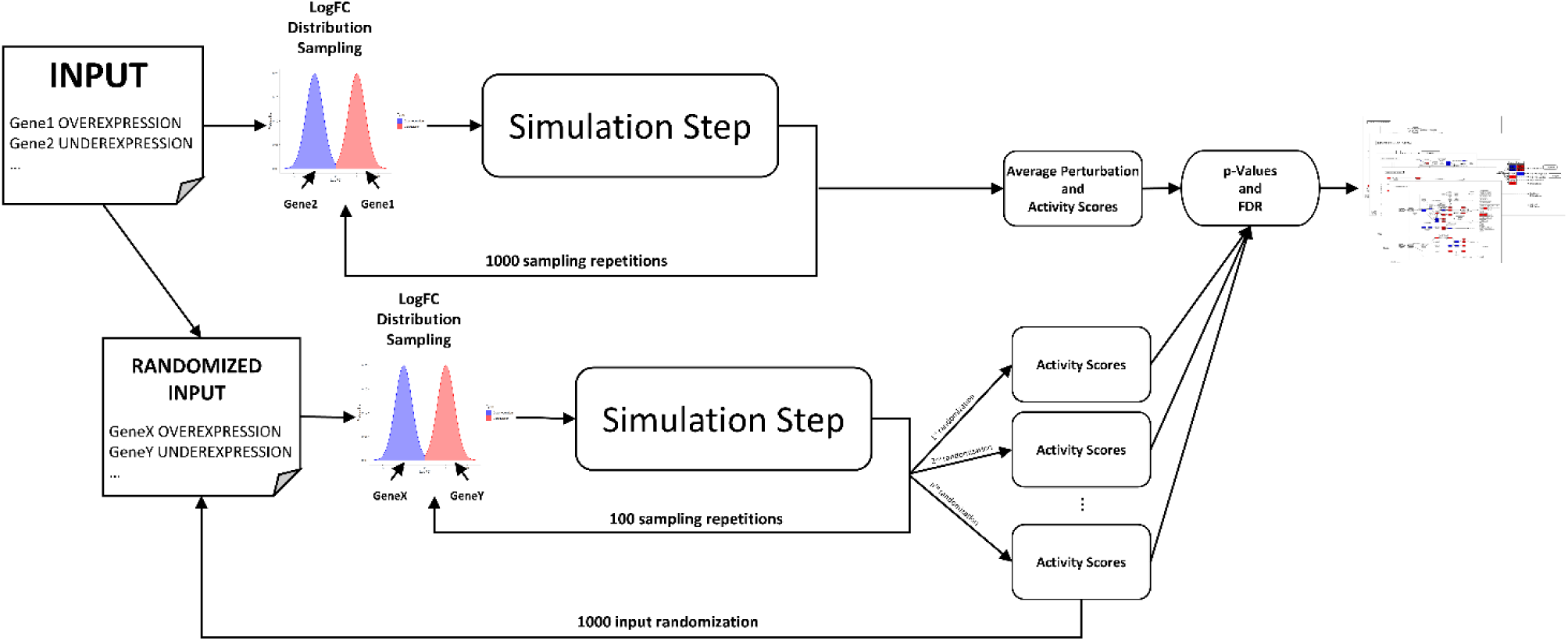
Description of the PHENSIM algorithm. First, the user provides a set of genes and the type of alteration (over-/under-expression). Then, synthetic LogFCs are generated, and a simulation step is performed. This is repeated 1000 times to compute the *Activity Scores*. Next, user input is randomized, and 100 synthetic LogFC are generated to estimate *Activity Scores* using the simulation step. This input randomization is repeated 1000 times for greater precision. Finally, p-values are computed, and the False Discovery Rate is estimated using the q-value methodology.

PHENSIM is implemented as a Java application for easy deployment on multiple operating systems. The source code is included in the MITHrIL platform and available at https://github.com/alaimos/mithril-standalone/tree/mithril-2.2. A web application is also available at https://phensim.atlas.dmi.unict.it/. All experimental data and source codes generated or analyzed during this study are available at https://github.com/alaimos/phensim.

### Synthetic LogFC generation

PHENSIM relies on MITHrIL perturbation analysis to compute the state of a node in the KEGG meta-pathway. Starting from LogFCs, MITHrIL propagates them through the network to estimate node and pathway perturbation. Hence a critical step in the PHENSIM simulator is the generation of Synthetic LogFCs.

By analyzing experimental data from “The Cancer Genome Atlas (TCGA),” we infer the space of feasible LogFCs. First, we got all cancer and control samples of TCGA to compute LogFCs of each gene for each cancer sample. With these data, we then fit two normal distributions for positive and negative LogFCs, respectively. This analysis produced two normal distributions with mean 5 for up-regulation (−5 for down-regulation) and standard deviation 2.

Synthetic LogFCs are estimated by sampling the two distributions. More precisely, let *x* ∈ {–1,0,1} be an input value, where –1 represents downregulation, +1 upregulation, and 0 no expression. At each simulation step, we generate a standard gaussian pseudorandom number, *r*_*𝒩*_(*x*), by using the polar method [31]. Synthetic LogFCs are then computed as:

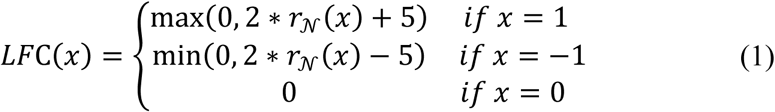

### PHENSIM Simulation Step

Let the KEGG meta-pathway be defined as a graph *G*(*V, E*) where *V* = {*V*_1_, *V*_2_, …, *V*_*m*_} is the set of all biological elements (genes, metabolites, miRNAs), and *E* ⊂ *V* × *V* is the set of activating or inhibiting interactions. Moreover, without loss of generality, we define PHENSIM input *𝒥* = {*V*_1_ = *ν*_1_, …, *V*_*n*_ = *ν*_*n*_} where *ν*_*k*_ ∈ {1,0, –1} for 1 ≤ *k* ≤ *n*, and *n* ≤ *m*. As previously described, we represent downregulation with –1, upregulation with +1, and no expression with 0.

To compute the activity of a biological element, each node *V*_*i*_ is considered as a discrete random variable that can assume three possible values: activated (1), inhibited (−1), or unchanged (0).

Given the input, the probability distribution of each variable is unknown. Therefore, we try to estimate it by generating synthetic LogFCs, which are then employed by MITHrIL perturbation analysis. Indeed, MITHrIL perturbation reflects the expected gene expression change when an alteration (expressed in terms of LogFC) is applied to a set of elements in the meta-pathway. Therefore, we collect these details to estimate a probability distribution empirically.

More in detail, given an input *𝒥*, at each step *t* of the simulation, we compute a set of LogFCs, Δ*E*_*𝒥*_(*k, t*) *for* 1 ≤ *k* ≤ *m*, where:

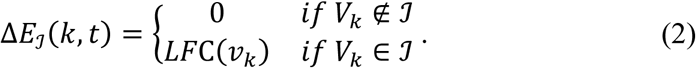

Next, for each node 0 ≤ *i* ≤ *m*, we estimate perturbation at step *t* as:

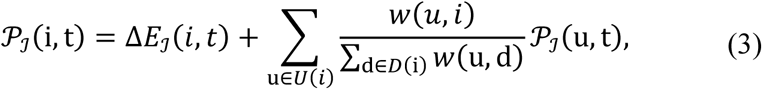

where *U*(*k*) and *D*(*k*) are the set of upstream and downstream nodes of *V*_*k*_, respectively, and *w*(*j, k*) is a weight reflecting the type of interaction between nodes *V*_*j*_ and *V*_*k*_. In PHENSIM, we use *w*(*j, k*) = 1 for all activating interactions, *w*(*j, k*) = –1 for all inhibiting ones. Finally, perturbations are returned for the computation of the *Activity Scores*.

### Activity Score Computation

Given the input *𝒥, PHENSIM* summarizes the activity of a node *V*_*i*_ in an A*ctivity Score, 𝒜*_*𝒥*_(*i*). The function of the *AS* is twofold. The sign gives the type of predicted effect: positive for activation, negative for inhibition. The value is the log-likelihood that such a result will occur. Therefore, to determine its value, we need to estimate the probability distribution of each node. To this end, we repeat the simulation step *𝒯* times to compute a set of perturbations *𝒫*_*𝒥*_(*i*) = {*𝒫*_*𝒥*_(*i, t*) *where* 1 ≤ *t* ≤ *𝒯*} for each node *V*_*i*_ of the graph.

Since the perturbation is negative for downregulation, positive for upregulation, and 0 for no alteration, we can use the sign function to determine node state. Therefore by counting the number of times each state appear during the simulation, we can empirically estimate the probability Pr(*V*_*i*_ = *ν*_*i*_|*𝒥*) for 1 ≤ *k* ≤ *m* as:

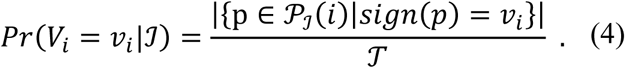

Finally, the activity score for a node *V*_*i*_ can be determined as:

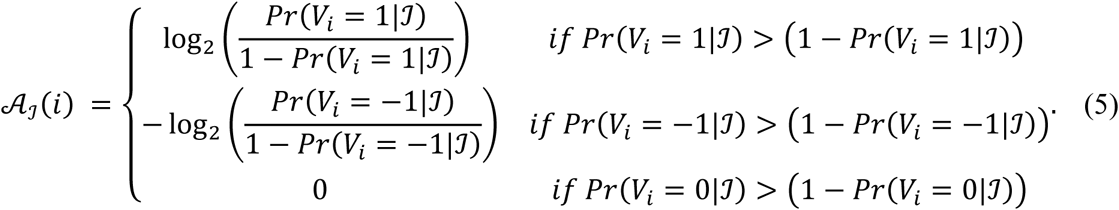

In all our experiments, we set *𝒯* = 1000 for the simulation step.

### Bootstrapping and Randomization

Together with the Activity Score, PHENSIM computes a p-value to establish which of the observed alterations are biologically relevant and not obtained by chance. Our idea is that a node is biologically relevant for the input if it is unlikely to observe a similar alteration when perturbing random nodes in the same way. We achieve this through a bootstrapping procedure together with input randomization. Given the *𝒥* = {*V*_1_ = *ν*_1_, …, *V*_*n*_ = *ν*_*n*_}, we compute *ℛ* random input set by taking arbitrary nodes from the KEGG meta-pathway. That is, for each randomization 1 ≤ *r* ≤ *ℛ*, we define a random input set 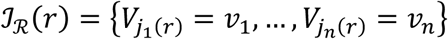 where 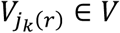 is a node of the meta-pathway chosen randomly in *V*. Next, for each input set, we compute synthetic LogFCs and run *𝒯* simulation steps to determine random *Activity Scores*, 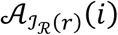. For the bootstrapping and randomization procedures, we set *ℛ* = 1000 and *𝒯* = 100.

### P-values computation and False Discovery Rate

PHENSIM p-value is empirically computed using the results from all simulations. Let *𝒜*_*𝒥*_(*i*) be the *Activity Score* computed for node 1 ≤ *i* ≤ *m* in the input simulation, and 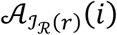 be the random *Activity Score* computed for an input randomization 1 ≤ *r* ≤ *ℛ*. We can say that a node alteration is not biologically relevant for the input if its probability is more significant than what might happen by chance. Therefore, if 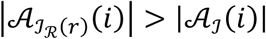 for most cases, we can say that the alteration is not specific for the simulated phenomena. We can synthesize this by using an empirically computed p-value as:

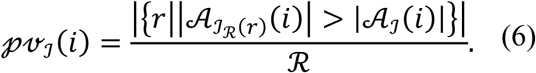

All p-values are then corrected for multiple hypotheses using the q-value approach and given as output together with the *Activity Score* and *Average Perturbation*.

### Dealing with dependent nodes

Equation 1 implies that all input nodes are altered independently from one another. However, we might want to simulate the case where two or more nodes are dependent. Since we do not always know how this dependency might alter the LogFC distribution, we can employ a simplified solution to address this. Indeed, we can modify the meta-pathway avoiding any changes to equation 1.

Let *𝒥* be the input and 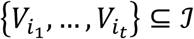 the dependent nodes, where 1 ≤ *i*_*k*_ ≤ *n* and 1 ≤ *k* ≤ *t* ≤ *n*. We can create a novel node *V*^*^ in the KEGG meta-pathway. Then, each edge connecting 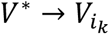 is built, and its weight is assigned as 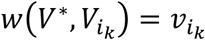, where 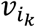 is the direction of the deregulation we wish to simulate. Therefore, we can build a new input set *𝒥*^*^, where all nodes are independent, as:

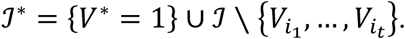

This new set can be used to approximate synthetic LogFC, taking dependencies into account, without estimating how such dependencies alter Log-Fold-Changes distribution. A detailed graphical representation of the process is depicted in figure S7.

### Experimental Procedure and Benchmarking

To assess PHENSIM prediction reliability, we built a benchmark based on data published in the GEO [32] database. More in detail, we want to determine how much PHENSIM can correctly predict the biological outcomes of the up-/down-regulation of a gene in a cell line through comparisons with expression data collected before and after the alteration. Therefore, we gathered 22 GEO series with such characteristics. Since different genes or cell lines could have been included in a single series, we obtained 50 case/control sample sets related to a specific gene/cell line combination. The details are shown in Supplementary Table S3. Each sample set was then divided into two categories, which were analyzed differently: (i) samples whose altered gene is present in the meta-pathway (called DS1), and (ii) samples whose altered gene is not in the meta-pathway (called DS2). For DS1, we directly simulated the alteration of the gene using PHENSIM. For DS2, we simulated the alteration of the differentially expressed genes (DEGs) computed between cases and controls. The rationale behind this choice is that DEGs somehow represent the effect of the source alteration.

For each dataset, non-expressed genes were identified according to the experiment type: Microarray or Sequencing. For sequencing, we chose all genes with an average count of less than 10. For microarrays, we selected all genes exhibiting an average expression less than the 10^th^ percentile. DEGs were computed using Limma [33] with a p-value threshold of 0.05 and a LogFC threshold of 0.6.

Each sample set was simulated as described above. Then, we compared the predicted perturbation (up/down-regulation) with LogFC computed on the expression data. All genes showing an absolute LogFC lower than 0.6 were considered as non-altered. Finally, we assessed the results in terms of accuracy (the number of correctly predicted genes divided by the total number of genes). Furthermore, since accuracy can be influenced by class imbalance, we chose to compute Positive Predictive Value (PPV), Sensitivity and Specificity for genes showing altered expression, and PPV and False Negative Rate (FNR) for the other genes. Indeed, for altered genes, PPV and Sensitivity represent the simulator’s ability to identify up-regulated genes (True Positives) correctly. In contrast, Specificity is the ability to identify down-regulated ones (True Negative). For non-altered genes, PPV is the rate of correctly identified genes with respect to all non-altered ones, while FNR is the rate of non-altered genes that were wrongly predicted as altered.

To compare performances with BioNSi, we ran the same simulations and computed the same metrics on the results. BioNSi requires an expression (in the range 0-9) for each gene and tracks how it changes until a steady-state is reached. Therefore, a gene is up-/down-regulated if the simulated expression increases/decreases between the initial and the final state, respectively. If no change is observed, the gene is not perturbed. To run the simulation, we loaded the meta-pathway and set all genes’ expression levels to 5. Next, we gave expression 9 for up-regulated gene, and 1 for down-regulated ones.

All raw data, input files, and other source codes are available for download at https://github.com/alaimos/phensim/tree/master/Benchmark.

## Author contributions

SA, AF, and AP conceived the work. AF and AP coordinated the research. BM participated in the evaluation of the model and the improvement of the reliability of the results. SA, ALF, GPM, BM, AP, and VR wrote the paper. SA developed the system. SA and VR build the benchmark. SA, ALF, GPM, and VR run the computational experiments. OBS and PNT provided biological insights and data. All authors analyzed the data and reviewed the final version of the manuscript.

