## Supplementary Materials for "PHENSIM: Phenotype Simulator"

### Supplementary Results

*Simulation #1: Anti-cancer effects of metformin*

Metformin is an agent for treatment of type 2 diabetes [1-4]. It inhibits glucose production in liver and increases insulin sensitivity in the peripheral tissues, resulting in elevated glucose uptake and consumption by skeletal muscle and adipose tissues. Metformin treatment reduces insulin secretion by β-pancreatic cells. The key molecule that performs these functions is AMP-activated protein kinase (AMPK), a serine-threonine kinase regulating cellular energy metabolism.

Several evidences have indicated that metformin possesses anti-cancer effects in various cancer types, especially in diabetic patients, in both direct and indirect manner [1-3]. Indeed, metformin directly activates the LBK1-AMPK signaling pathway [3]. Metformin is known to uncouple the electron transport chain in the mitochondria by targeting Complex I ([1, 2, 5]), leading to impaired mitochondrial function, decreased adenosine triphosphate (ATP) synthesis, and elevated cellular AMP/ATP ratio [1, 3]. Increased binding of AMP to AMPK activates AMPK by inducing phosphorylation of its catalytic subunit at residue Thr172 by liver kinase B1 (LKB1), a tumor suppressor and a regulator of cellular energy status [2, 3]. Binding of AMP to AMPK also prevents dephosphorylation of AMPK Thr172 by protein phosphatases. LKB1-activated AMPK phosphorylates and activates the tumor suppressor Tuberous Sclerosis Complex 1 and 2 (TSC1/2), which negatively regulates the activity of mammalian target of rapamycin (mTOR), which is upregulated in most cancer cells and causes tumor proliferation and cell growth, by inhibiting Ras homolog enriched in brain (Rheb) [1, 3]. mTOR is a critical mediator of the phosphatidylinositol-3-kinase/protein kinase B/Akt (PI3K/PKB/Akt) signaling pathway, which is one of the most frequently deregulated molecular networks in human cancer [3].

Metformin-activated AMPK inhibits mTOR and reduces the phosphorylation of its downstream targets, the eukaryotic initiation factor 4E-binding proteins (4EBPs) and ribosomal S6 kinases (S6Ks), leading to an inhibition of global protein synthesis, cell cycle progression, cell proliferation and angiogenesis [3]. Moreover, AMPK has been reported to suppress mTOR signaling pathway independent of TSC2 via phosphorylation of mTOR binding protein Raptor.

Metformin has been shown to cause a G0/G1 cell cycle arrest by decreasing the expression of cyclin D1 [2].

Metformin-induced AMPK activation has been shown to phosphorylate insulin receptor substrate-1 (IRS-1) at Ser-794 which results in decreased recruitment of the p85 subunit of phosphoinositide-3-kinase (PI3K), thus, impairing the insulin-like growth factor (IGF)-stimulated PI3K/protein kinase B/ mammalian target of rapamycin complex 1 (PI3K/Akt/mTORC1) signaling pathway.

Metformin also inhibits the crosstalk between G-protein-coupled receptors (GPCR) and insulin/IGF1 receptors signaling, resulting in the inhibition of mTORC1 and in reduction of cellular proliferation [1, 2].

Metformin induces nuclear degradation and decreased expression of Sp proteins, transcription factors for genes involved in cell proliferation (cyclin D1), metabolism (FAS), apoptosis (B-cell lymphoma 2, bcl-2, and survivin) and angiogenesis (vascular endothelial growth factor, VEGF, and its receptor VEGFR1) [2, 3].

The indirect mechanism of metformin in anti-cancer function is related to its ability to lower insulin and insulin-like growth factor 1 (IGF-1) [3].

Metformin disrupts insulin and IGF-1 signaling pathways by reducing insulin and IGF-1 levels, reducing total IGF-1 receptor and IR levels, and downregulating IGF-1 receptor and IR gene expression [6].

In parallel with this, metformin also downregulates the MAPK (mitogen-activated protein kinase) pathway, NF-$\kappa$B (nuclear factor kappa B) signaling [5, 7], glycolysis, and the TCA (tricarboxylic acid) cycle [2, 3, 6].

Based on these details, we run a PHENSIM simulation of the simultaneous upregulation of LKB1 and the downregulation of insulin (Ins), IGF1, and GPD1 [8].

As expected, PHENSIM returned significant downregulation of Insulin (pathway activity score = -8.7121, p-value 0.072) and mTOR signaling (pathway activity score = -8.7121, p-value 0.075).

All isoforms of PI3Ks (phosphoinositide 3-kinases) and AKT (serine/threonine protein kinase Akt) as well as the metabolite PIP3 (phosphatidylinositol (3,4,5)-trisphosphate) were also downregulated (PIK3CA activity score = -4.8203, p-value 0.021; AKT1 activity score = -4.2546 p-value 0.04).

Although the negative regulation of mTOR (activity score = -3.9992; p-value = 0.04) should activate the *repressor of translation initiation* 4EBP, the simulation returns no activity score for this node. However, a low positive perturbation for 4EBP can be observed (perturbation = 0.00052). PHENSIM also predicted the inhibition of downstream nodes involved in protein synthesis such as S6Ks (fig. S1a) (S6K-alpha3 activity score = -3.4095, p-value = 0.027; S6K-alpha2 = -1.3129, p-value= 0.013).

We can also predict a low perturbation for TCA cycle pathway (pathway perturbation = -0.0000177).

MAPK and NF-$\kappa$B (nuclear factor kappa B) signaling were predicted as downregulated (MAPK pathway activity score = -8.7121, p-value 0.074; NF-$\kappa$B activity score = -4.8203, p-value = 0.111). For these two pathways, several downregulated enzymes and metabolites were predicted, in full agreement with data from literature [6] (fig. S1b and tab. S2).

Finally, in accordance with literature, PHENSIM also predicted weak changes in cytokine gene expression as it can be seen from average nodes perturbations (IL6 perturbation = -0.00000135; IFN-α perturbation = -0.000001238; IL8 perturbation = - 0.00000227; IL17 perturbation = -0.000000382; TNF-alpha perturbation= -0.0006175) [6].

*Simulation #2: Everolimus (RAD001) and breast cancer*

Everolimus (RAD001, Afinitor®), an analog of rapamycin, has shown immunosuppressive and anticancer activities [9-12]. It is currently approved for treatment of various types of cancer including metastatic breast cancer [13-15]. Everolimus has a growth inhibitory activity against tumor cells and can retard tumor growth through direct mechanisms against both the tumor cell and the solid tumor stroma components [11].

Everolimus inhibits *mammalian target of rapamycin* (mTOR), to prevent the downstream signaling required for cell cycle progression, cell growth, and proliferation [9-11, 16-18].

In mammalian cells, mTOR exists in two complexes, mTORC1 and mTORC2 [14, 15, 17, 19], which are differentially regulated and have distinct substrate specificities [14]. mTORC2 signaling, is lower in breast tumors compared to normal breast tissue. This could suggest that mTORC1 signaling is more oncogenic than mTORC2 [20].

mTORC1 promotes protein synthesis by: (a) stimulating ribosome biogenesis via phosphorylation and inhibition of the RNA Polymerase III repressor MAF1 [21]; (b) phosphorylating the p70S6K and 4EBP1 and modulating the activity of their downstream targets [22, 23]; (c) by regulating nucleocytoplasmic RNA transport [21, 22]. In addition, mTORC1 stimulates pyrimidine biosynthesis and lipid biosynthesis [22, 23]. mTORC1 phosphorylates ULK1 (unc-51 like autophagy activating kinase 1) and DAP (death-associated protein) inhibiting autophagy [20, 24].

Finally, upregulation of mTOR signaling can promote tumor growth and progression through several mechanisms including the promotion of growth factor receptor signaling, angiogenesis, glycolytic metabolism, lipid metabolism, cancer cell migration, and suppression of autophagy [14, 15].

All these functions of mTORC1 are reversed by Everolimus and other mTORC1 inhibitors [10, 13] (fig. S2).

Everolimus binds with high affinity to its intracellular receptor, the FKBP12, a protein belonging to the immunophilin family. The everolimus–FKBP12 complex binds mTOR when it is associated with RAPTOR and mLST8 to form mTORC1 complex, resulting in decreased interaction between mTOR and RAPTOR, which could inhibit the phosphorylation and activation of the major mTORC1 downstream targets [12, 14-16, 19, 20].

Here we wanted to simulate the inhibition of mTORC1. Unfortunately, simulating mTORC1 inhibition was not feasible because KEGG does not distinguish the mTOR node in mTORC1 from the one included in mTORC2. To overcome such limitation, we have set the downregulation of p70S6K (p70S6Ka and p70S6Kb) and 4EBP and the upregulation of ULK1/2 because these are the well-known downstream targets of mTORC1. Then we uploaded a list of unexpressed genes in breast tissue to simulate the effects of the drug on such a tissue. Our simulation predicted that factors associated with RNA transport would be downregulated, while factors involved in autophagy would be upregulated. The simulation showed that RNA transport signaling pathway exhibits a low activity scores (activity score = -4.8203; p-value = 0.013) (fig S3a). Furthermore, we could predict several downregulated factors involved in RNA transport and protein synthesis, such as eukaryotic translation initiation factor 4A, 4B and ribosomal proteins S6Ks, p70-S6K and p70S6Kb (eIF4A activity score = - -4.8203; eIF4B activity score = -4.8203; p70-S6K activity score = -4.8203; p70S6Kb activity score = -4.8203; p-value for all nodes < 0.01). PHENSIM also predicts the 4EBP1 inhibition (activity score = -4.8203; p-value = 0.003) and consequently the upregulation of eIF4E (activity score = 4.8203; p-value = 0.003) (fig S3b).

Upregulation of the autophagy (activity score = 2.0687, p-value 0.254) was a consequence of alterations in ULK1/2 phosphorylation levels. However, PHENSIM failed in predicting the deregulation of p21 (cyclin-dependent kinase inhibitor 1), cyclin D and NF-kB [13]. This limitation is probably due to the presence of a single node for mTORC1 and mTORC2.

*Simulation #3: effects of exosomal vesicles on hematopoietic stem/progenitor cells (HSPCs) in the bone marrow (BM)*

The functional relevance of cancer-derived exosomes to tumor growth, metastasis and treatment response has become increasingly evident [25, 26]. Exosomes derived from AML blasts contain complex cargoes which function via paracrine mechanisms to modulate the properties of both the tumor cells themselves and the BM niche. Several microRNAs have been shown to be selectively incorporated in these exosomes, including miR-150 and miR-155 [27, 28]. One of the targets of these microRNAs is the transcription factor c-MYB, which is downregulated in tumor cells exposed to the exosomes [28]. Additional targets include c-KIT, DNMT1, Lymphoid Cell Helicase (HELLS), PAICS, an enzyme involved in purine biosynthesis, TAB2, and others. The downregulation of these molecules compromises hematopoiesis via stroma-independent mechanisms. However the cargo of AML cell-derived exosomes also targets mesenchymal stromal progenitors, inhibiting/reducing the expression of hematopoietic stem cell supporting factors such as CXCL12 (C-X-C motif ligand 12), KITL (c-Kit ligand), IL-17 and IGF1 and interfering with both hematopoiesis and osteogenesis [25] (fig. S4). Moreover AML-derived exosomes increase expression of genes supporting AML growth (DKK1, IL-6, CCL3).

To determine whether PHENSIM can make the correct predictions in this model, we ran a simulation for the uptake of the eight most representative miRNAs (miR-150, -155, -146a, -191, -221, -99b, -1246 and let-7a) included in AML-derived exosomes by hematopoietic stem cells [27].

The simulation predicts an inhibition of osteoclast differentiation (activity score = -8.7121, p-value=0.103) and cytokine-cytokine receptor interaction pathways (activity score = -4.8203, p-value=0.086) (fig. S5a-b).

In agreement with the literature, some genes involved in modulation of normal hematopoiesis, like CXCL12 (activity score = -4.5951, p-value= 0.003) and the receptor IGF1R (activity score = -4.8203, p-value= 0.022), but not IGF1, were downregulated [25]. Similarly, c-MYB, which is involved in

HSPC differentiation and proliferation, was also downregulated [28] (activity score = -4.7015; p-value = 0.009) (fig. S5c). However, PHENSIM failed to predict the upregulation of DKK, IL6 and CCL3, and the downregulation of KITL and IL17 (activity score = 0) [25].

*Simulation #4: testing TNFα/siTPL2-dependent synthetic lethality on a subset of human cancer cell lines*

TNFα (tumor necrosis factor alpha), a type II transmembrane protein, is a member of the tumor necrosis factor cytokine superfamily and has an important role in innate immunity and inflammation. Although it has the potential to induce cell death, most cells are protected by a variety of mechanisms. In a recent paper, Serebrennikova et al. [29] showed that one of the checkpoints of TNFα-induced cell death is TPL2 (MAP3K8), a MAP3 kinase that is known to have an important role in immunity, inflammation, and oncogenesis. The knockdown of TPL2 resulted in the downregulation of miR-21 and the upregulation of its target CASP8 (caspase-8). This combined with the downregulation of the caspase-8 inhibitor cFLIP (FADD-like IL-1β-converting enzyme inhibitory protein), resulted in the activation of caspase-8 by TNFα and the initiation of apoptosis (fig. 4). The activation of caspase-8 also promotes the activation of the mitochondrial pathway of apoptosis, although some molecules such as BIML (Bcl-2-like protein 11, isoform L), which are also involved in the activation of the mitochondrial pathway, may be activated via caspase-8-independent mechanisms. An important upstream regulator of this pathway is NF-κB. The activation of ERK (MAPK1/2), JNK (c-Jun Nterminal kinase) and p38MAPK, the activation of AKT and the phosphorylation of GSK3 (glycogen synthase kinase 3) at Ser9/21 are also inhibited by the knockdown of TPL2. However, their inhibition does not appear to have a role in the initiation of TNFα/siTPL2-induced apoptosis. It is worth noting that the activation of the apoptotic (caspase-8-dependent) pathway in TNFα/siTPL2 treated cells was observed in some, but not all cancer cell lines, suggesting that correct prediction will depend on whether the data analyzed by PHENSIM are derived from sensitive or resistant cells.

To launch the simulation, we set TPL2 and miR-21-5p as downregulated and TNFα as upregulated. Since our goal was to simulate the outcome of such treatment in six cell lines, i.e. HeLa, HCT116, U2-OS, CaCo-2, RKO and SW480, we launched six different simulation. Each simulation had a different list of non-expressed genes, one for each cell-line.

Among these tumor cell lines, only HeLa, HCT116, U2-OS were sensitive to treatment with TNFα/siTPL2. At the end of the computations, PHENSIM was able to predict the upregulation of caspase-8 for any of the six cell lines. However, PHENSIM could not predict the downregulation of cFLIP in the resistant cell lines. This could be the result of missing information in KEGG pathways. Also, PHENSIM did not predict any activity score for MLC1 (Mcl-1 apoptosis regulator) and XIAP (X-linked inhibitor of apoptosis) nodes, although the system predicts a low negative perturbation for both genes in all cell lines except for HCT116 where the opposite happens.

PHENSIM predicted the upregulation of the apoptosis inhibitors BCL2 and BCL-XL in all sensitive cell lines (only for HCT116 the activity score is 0 but the average node perturbation is still positive). BCL2 is upregulated also in RKO and SW480 cells. PHENSIM showed a negative perturbation of the inducer of mitochondrial apoptosis BAX only in U2-OS and HELA among the sensitives, and SW480 and SW480 as resistant cell lines (fig. S6a-d).

Although these results do not exactly reflect our expectations as there are discrepancies between the in vitro and in silico experiment done by PHENSIM, it was confirmed by results obtained by the previously mentioned experimental study [29], which suggested that the change in the expression of such molecules was due to the activation of feedback mechanisms. Interestingly, this result was obtained only for four out of six cancer cell lines, of which three were sensitive (HeLa, HCT116 and U2-OS) and one was resistant (CaCo-2).

In addition, phosphorylated ERK, MEK, JNK and p38 activity were strongly downregulated for all of the six cell lines, but not cIAP2 (baculoviral IAP repeat containing 2) which has activity score 0 and a weak negative perturbation, in HTC116, HeLa, U2OS, SW480 and CaCo-2 cells, as confirmed by the experimental data (fig S6a-b).

### Supplementary Figures

a)

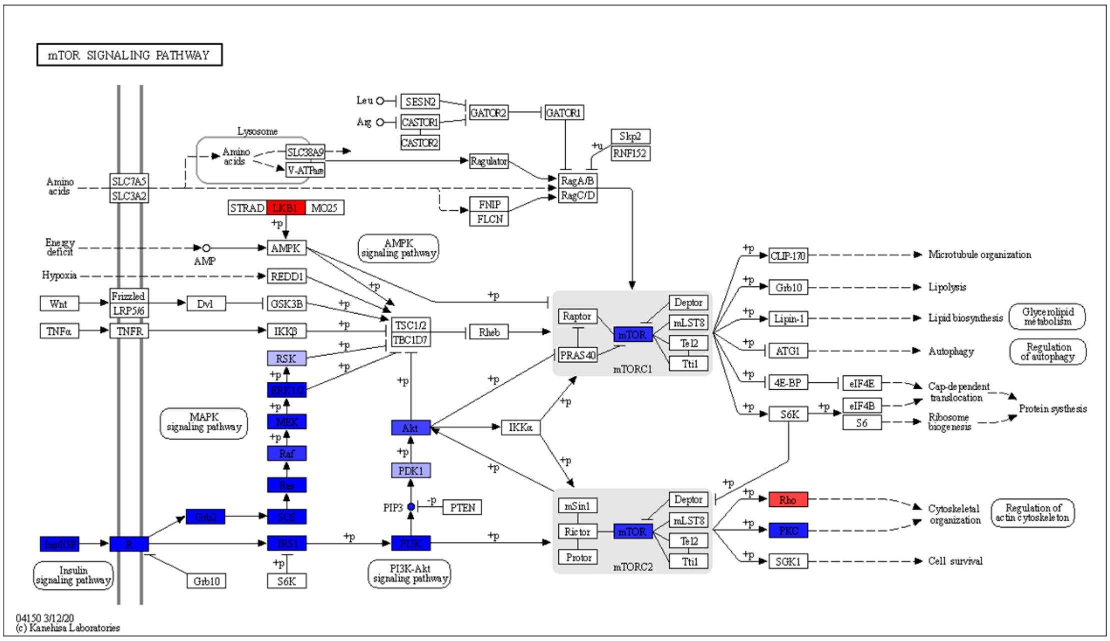


b)

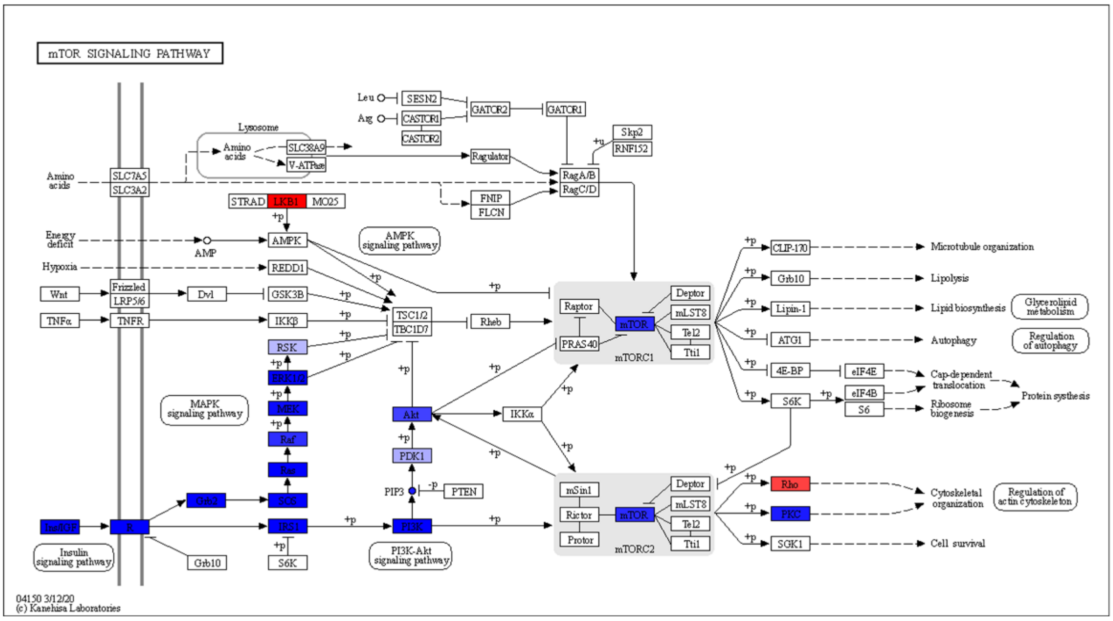


**Figure S1.** Anti-cancer effects of metformin predicted by PHENSIM. The simulation was launched by assuming downregulation of INS and IGF-1, and upregulation of LKB1. In **(a)** are shown predictions related to the mTOR signaling. In **(b)** are shown predictions related to a subset of nodes belonging to the MAPK and NF-kB signaling and involved in the TNF signaling pathway. Downregulated nodes are colored in blue. Upregulated nodes are colored in red.

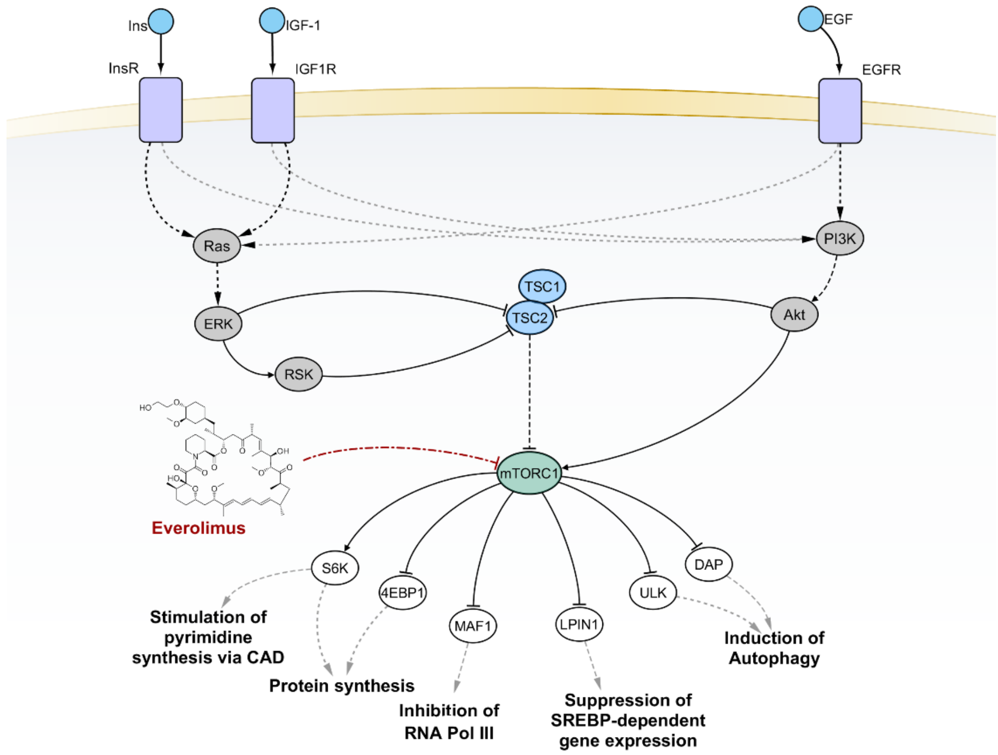


**Figure S2.** mTORC1 and its downstream signaling pathways. Black solid edges represent direct interaction between first neighbor nodes. Dashed edges represent indirect interactions between nodes. Red dot-dashed edges evidence scientifically validated interactions considered for PHENSIM prediction.

a)

| ID | NAME | ACTIVITY SCORE | P-VALUE |
|---|---|---|---|
| path:hsa04910 | Insulin signaling pathway | -7.9983 | 0.1080 |
| path:hsa04151 | PI3K-Akt signaling pathway | -7.9983 | 0.1210 |
| path:hsa04371 | Apelin signaling pathway | -7.9983 | 0.1060 |
| path:hsa04714 | Thermogenesis | -7.9983 | 0.1020 |
| path:hsa04152 | AMPK signaling pathway | -7.9983 | 0.1070 |
| path:hsa04724 | Glutamatergic synapse | -7.6768 | 0.1390 |
| path:hsa04072 | Phospholipase D signaling pathway | -7.5718 | 0.1070 |
| path:hsa04935 | Growth hormone synthesis, secretion and action | -7.3467 | 0.1150 |
| path:hsa04723 | Retrograde endocannabinoid signaling | -7.2503 | 0.1390 |
| path:hsa04920 | Adipocytokine signaling pathway | -6.6784 | 0.0890 |
| path:hsa04919 | Thyroid hormone signaling pathway | -6.5952 | 0.1160 |
| path:hsa04150 | mTOR signaling pathway | -6.3366 | 0.1110 |
| path:hsa04211 | Longevity regulating pathway | -6.3035 | 0.1140 |
| path:hsa04066 | HIF-1 signaling pathway | -5.2604 | 0.1120 |
| path:hsa04270 | Vascular smooth muscle contraction | -5.2167 | 0.1600 |
| path:hsa04659 | Th17 cell differentiation | -5.0614 | 0.1000 |
| path:hsa04350 | TGF-beta signaling pathway | -4.8203 | 0.0910 |
| path:hsa00561 | Glycerolipid metabolism | -4.8203 | 0.0280 |
| path:hsa04666 | Fc gamma R-mediated phagocytosis | -4.8203 | 0.1110 |
| path:hsa04012 | ErbB signaling pathway | -4.8203 | 0.1020 |
| path:hsa04630 | JAK-STAT signaling pathway | -4.8203 | 0.1110 |
| path:hsa03320 | PPAR signaling pathway | -4.8203 | 0.0450 |
| path:hsa04514 | Cell adhesion molecules (CAMs) | -4.8203 | 0.0310 |
| path:hsa04218 | Cellular senescence | -4.8203 | 0.1230 |
| path:hsa04213 | Longevity regulating pathway - multiple species | -4.8203 | 0.1090 |
| path:hsa03013 | RNA transport | -4.8203 | 0.0130 |
| path:hsa04713 | Circadian entrainment | -4.7215 | 0.0930 |

b)

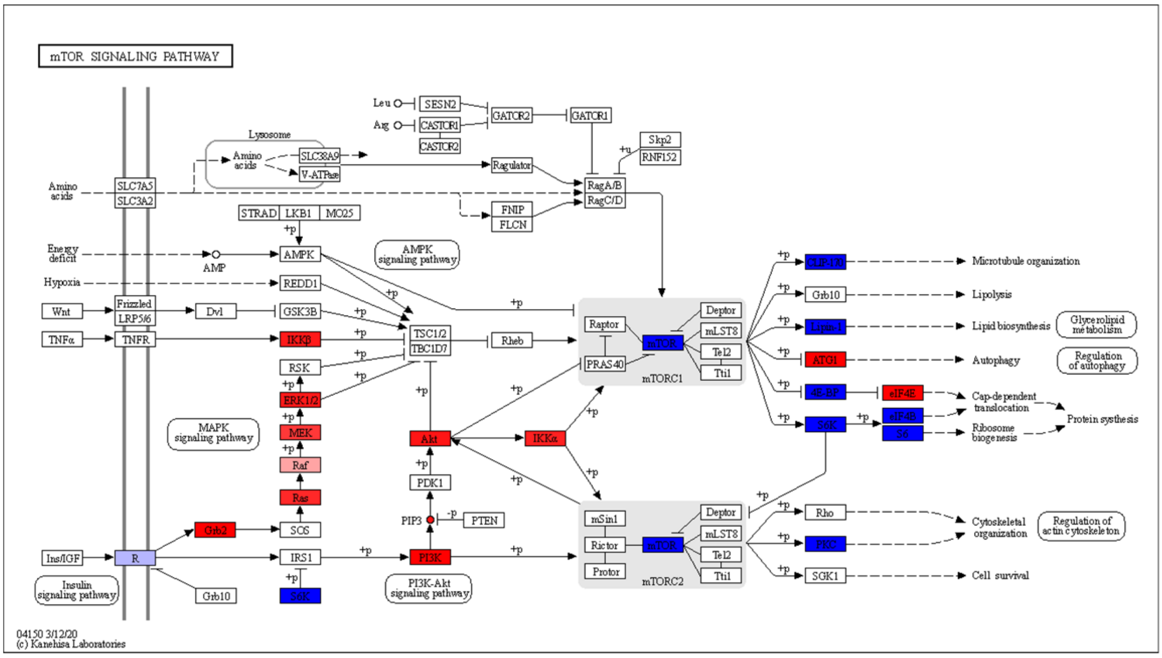


**Figure S3.** Prediction of perturbation on pathways in mammary tissue caused by everolimus. Figure **(a)** reports the top 10 list of negatively deregulated pathways, among which figured both the RNA transport and mTOR signaling pathways. In **(b)** are shown predictions related to the mTOR signaling. Downregulated nodes are colored in blue. Upregulated nodes are colored in red.

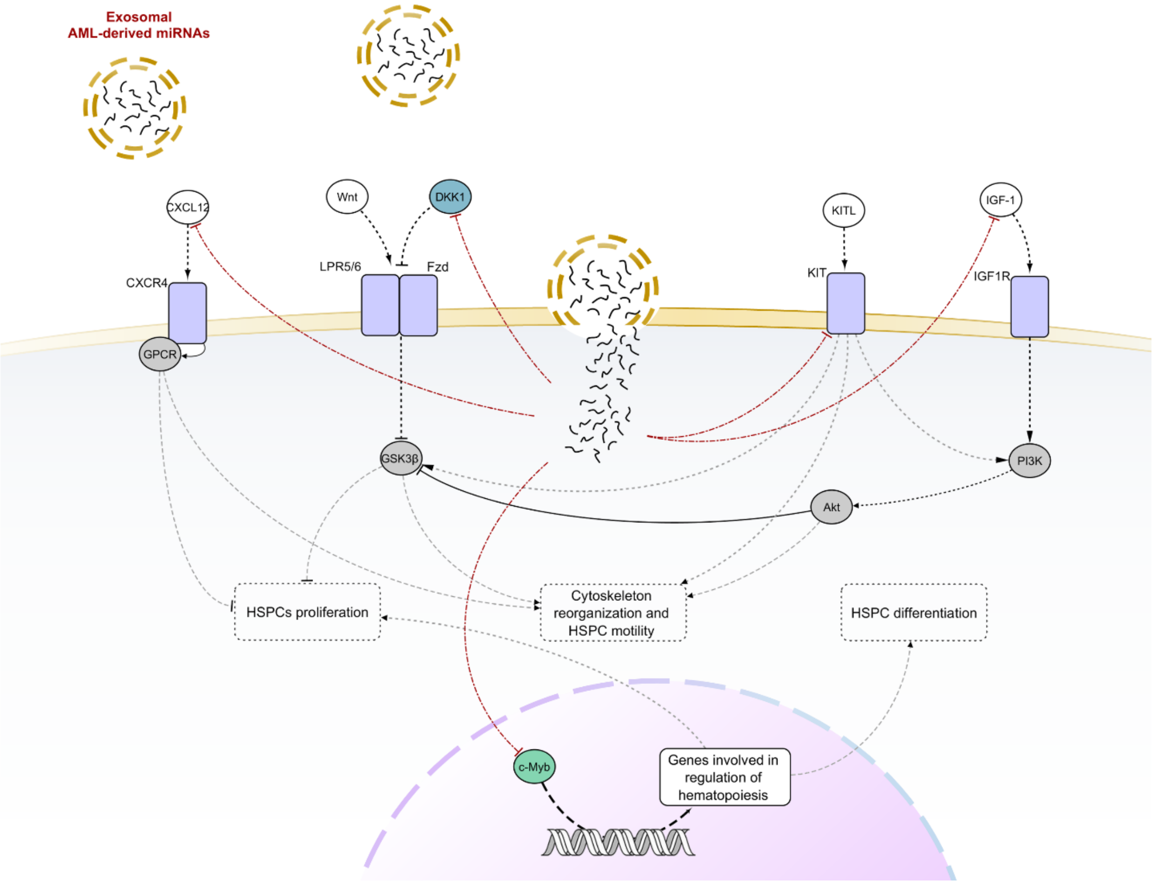


**Figure S4**. A reconstructed model showing cellular components involved in hematopoiesis and motility of HSPCs and their downregulation mediated by exosomal-miRNAs derived from AML cells. Black solid edges represent direct interaction between first neighbor nodes. Dashed edges represent indirect interactions between nodes. Red dot-dashed edges evidence scientifically validated interactions considered for PHENSIM prediction.

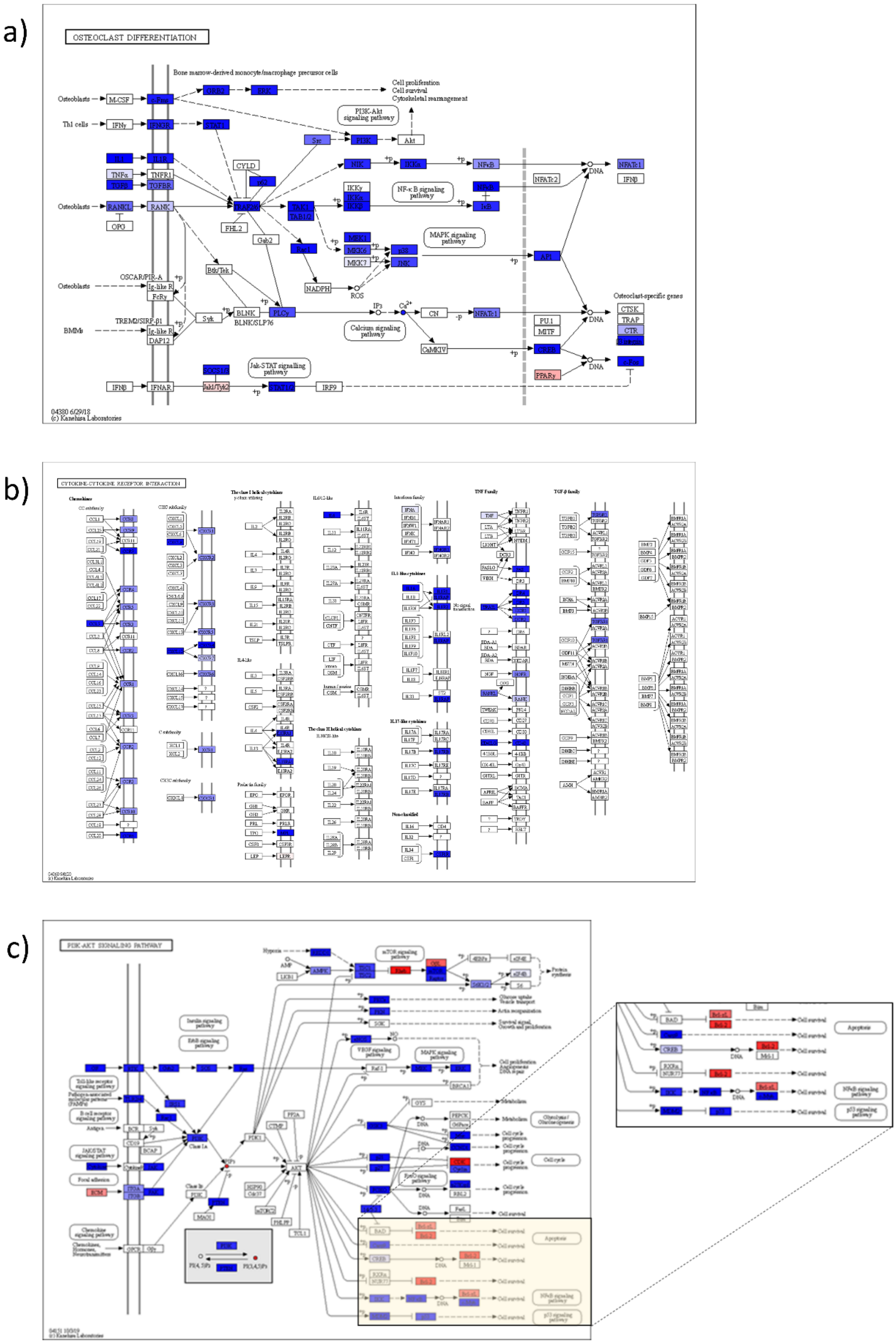


**Figure S5.** Prediction of perturbations caused by AML-derived exosomal miRNAs on recipient cells. Figures **(a)** and **(b)** show a wide deregulation of several nodes belonging to the Osteoclast differentiation pathway and Cytokine-cytokine receptor interaction pathway, respectively. In **(c)** is shown downregulation of c-Myb within the PI3K-Akt signaling pathway. Downregulated nodes are colored in blue. Upregulated nodes are colored in red.

a)

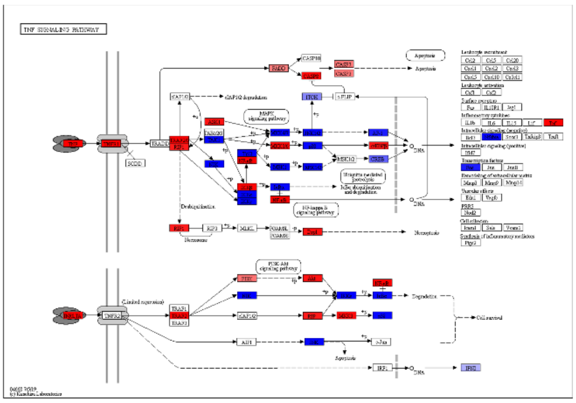

b)

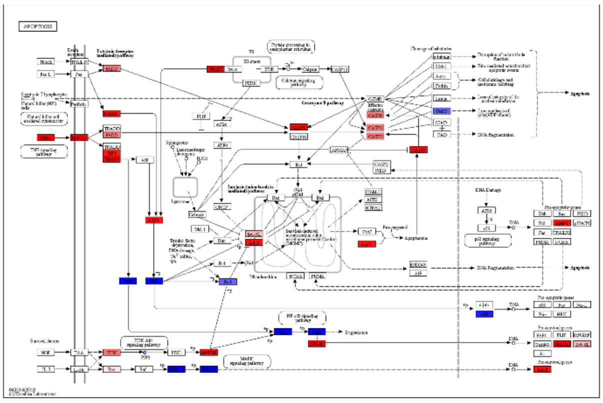

c)

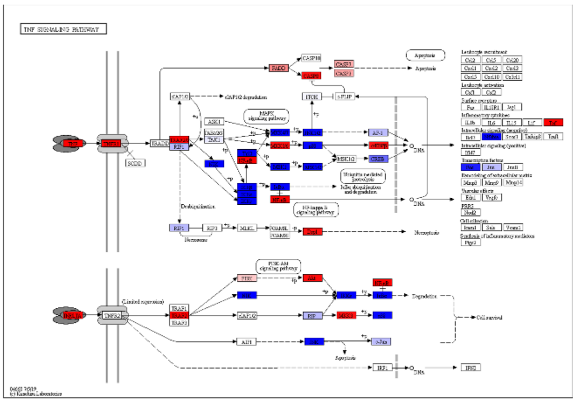

d)

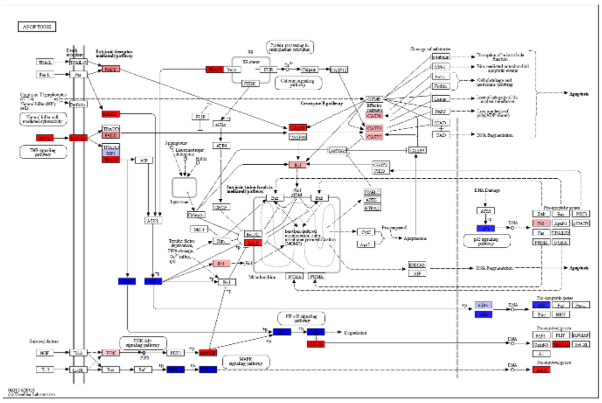

**Figure S6.** Effects of TPL2 KD and TNFα simulated by PHENSIM. In **(a)** and **(b)** are shown results obtained for TNF signaling and Apoptosis pathway respectively, in the context of HeLa cells (chosen as representative for sensitive cell lines). In **(c)** and **(d)** are shown results obtained for TNF signaling and Apoptosis pathway respectively, in the context of RKO cells (chosen as representative for resistant cell lines). Results for CaCo-2 cells, for which PHENSIM returned a pattern of deregulation like that of sensitive cell lines, are not shown. Downregulated nodes are colored in blue. Upregulated nodes are colored in red.

$$\mathcal{I} = \{V_1 = 1, V_2 = 1, V_3 = -1\} \qquad \mathcal{I}^* = \{V_1 = 1, V^* = 1\}$$

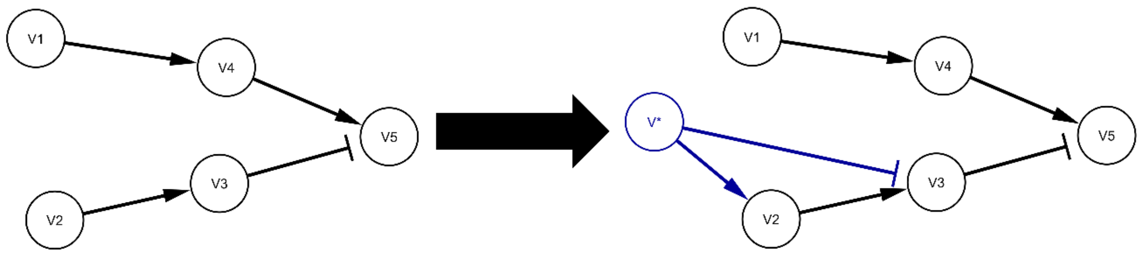


**Figure S7.** Manipulation of the meta-pathway when simulating dependent nodes. We wish to simulate upregulation of nodes $V_1$ and $V_2$ and downregulation of $V_3$. Since we know that the expression of $V_2$ and $V_3$ are dependent, we add a novel node $V^*$ which activates $V_2$ ($w(V^*, V_2) = 1$) and inhibits $V_3$ ($w(X^*, X_2) = -1$). Finally, we can run the simulation by upregulating both nodes $V_1$ and $V^*$.

### Supplementary Tables

| | Algorithm | Accuracy | Altered Genes | | | Non-altered Genes | |
|---|---|---|---|---|---|---|---|
| | | | PPV | Sensitivity | Specificity | PPV | FNR |
| **Datasets where input gene was in KEGG** | | | | | | | |
| | **PHENSIM** | 0.6647 | 0.5123 | 0.5191 | 0.9884 | 0.8920 | 0.2711 |
| | **BioNSi** | 0.0640 | 0.5075 | 0.2692 | 0.7925 | 0.8624 | 0.9970 |
| **Datasets where input gene was not in KEGG** | | | | | | | |
| | **PHENSIM** | 0.5977 | 0.7725 | 0.8421 | 0.9885 | 0.9000 | 0.4091 |
| | **BioNSi** | 0.0735 | 0.3283 | 0.1500 | 0.8052 | 0.4345 | 0.9968 |

**Table S1. Summary of the comparisons between PHENSIM and BioNSi**. We computed for both software accuracy, Positive Predictive Value (PPV), Sensitivity and Specificity for genes showing altered expression, and PPV and False Negative Rate (FNR) for the non-altered ones. All 50 sample sets used for the benchmark were categorized based on the gene presence in the KEGG meta-pathway.

| MAPK signaling pathway | | | |
|---|---|---|---|
| **Gene Name** | **Perturbation** | **Activity Score** | **p-value** |
| NF-kB, p50 | -0.000704156 | 0 | 0.564 |
| NFKB2, p52 | -4.01112E-05 | 0 | 0.921 |
| RAC1 | -0.00205679 | -2.14286335 | 0.05 |
| NRAS | -0.075572228 | -4.820281317 | 0.031 |
| MRAS | -0.019194153 | -4.595119651 | 0.013 |
| HRAS | -0.073178847 | -4.820281317 | 0.029 |
| R-Ras | -0.008108077 | -4.254598883 | 0.025 |
| RASGRP3, GRP3 | -0.007319367 | -4.119037051 | 0.016 |
| RRAS2 | -0.05580463 | -4.820281317 | 0.014 |
| KRAS | -0.073178847 | -4.820281317 | 0.03 |
| RASA1 | -0.003321088 | -3.178053781 | 0.006 |
| RAC1 | -0.00205679 | -2.14286335 | 0.05 |
| CDC42 | 0.001932396 | 1.982994337 | 0.029 |
| **NF-kappa B signaling pathway** | | | |
| **Gene Name** | **Perturbation** | **Activity Score** | **p-value** |
| NF-kB, p50 | -0.000704156 | 0 | 0.564 |
| NFKB2, p52 | -4.01112E-05 | 0 | 0.921 |
| IKBA | -0.000504171 | 0 | 0.743 |

**Table S2. Summary of the predictions for the MAPK and NF-$\kappa$B signaling pathways**. Here, we report the most important predictions made by PHENSIM for the MAPK and NF-$\kappa$B signaling pathways. For each pathway we report a set of relevant nodes together with their perturbation, activity score, and their p-value.
